# Microbial Metagenomic Approach Uncovers the First Rabbit Haemorrhagic Disease Virus genome in Sub-Saharan Africa

**DOI:** 10.1101/2020.11.19.390559

**Authors:** Anise N. Happi, Olusola A. Ogunsanya, Judith U. Oguzie, Paul E. Oluniyi, Alhaji S. Olono, Jonathan L. Heeney, Christian T. Happi

**Affiliations:** Department of Veterinary Pathology, Faculty of Veterinary Medicine, University of Ibadan, Ibadan, Nigeria; Department of Biological Sciences, Faculty of Natural Sciences, Redeemer’s University, Ede, Nigeria; African Centre of Excellence for Genomics of Infectious Diseases (ACEGID), Redeemer’s University, Ede, Osun State, Nigeria; Lab of Viral Zoonotics, Department of Veterinary medicine, University of Cambridge, Cambridge, UK

**Keywords:** Rabbit Haemorrhagic disease, Nigeria, RHDV2

## Abstract

Rabbit Haemorrhagic Disease (RHD) causes high morbidity and mortality in rabbits and hares. Here, we report the first genomic characterization of Rabbit Hemorrhagic Disease Virus (RHDV) from sub-Saharan Africa. While suspected, only a single PCR finding without sequence confirmation or characterization has been reported. Here, we used a microbial metagenomic approach to confirm and characterize pathogens causing the suspected outbreak of RHD in Ibadan, Nigeria. On the 25^th^ September 2020, the liver, spleen, and lung samples of five rabbits from an outbreak in 2 farms in Ibadan, Nigeria, were analyzed for the vp60 gene of RHDV by RT-PCR. Subsequently, Next Generation Sequencing on 1^st^ of October revealed one full and two partial RHDV2 genomes on both farms. Phylogenetic analysis showed close clustering with RHDV2 lineages from Europe, in particular, 98.6% similarity with RHDV2 in the Netherlands, and 99.1 to 100% identity with RHDV2 in Germany, suggesting potential importation from Europe. The detection of twelve unique mutations in RHDV2 sequences from the Ibadan outbreak compared to other RHDV2 sequences in the same clade suggests significant genetic diversity of the GI.2 strains in Nigeria. This highlights the need to further understand the genetic diversity of Lagoviruses to, inform vaccine development, and for accurate tracking, monitoring, and control of outbreaks in Africa.

## Introduction

Rabbit haemorrhagic disease (RHD) is a highly infectious and deadly viral haemorrhagic disease of rabbits. The disease is caused by Rabbit haemorrhagic disease virus (RHDV) a *Lagovirus* of the *Caliciviridae* family^1^. The virus causes a high morbidity and mortality rates, killing more than 90% of infected adult animals in 2-3 days following infection^2,3^. This disease causes economic losses to the rabbit meat and fur industry and great negative ecological impact in wild rabbit populations^3, 4^. RHD is among the diseases notifiable to the World Organization for Animal Health (OIE).

Transmission is via nasal, oral, conjunctival routes with possible vector-borne transmission reported^5^. The virus is also shed through excretions from infected animals^6^. The lesions of RHD include petechial hemorrhages in multiple organs as a result of virus-induced hyper-coagulopathy. The most severe form of these lesions appears in the liver, trachea, and lungs^7^. RHDV also promotes fatal hepatitis in adult rabbits^8^.

The RHDV genome is positive sense and single stranded RNA of approximately 7437 nucleotides in length^1^. There are two open reading frames (ORFs); ORF1 encoding the nonstructural proteins (RNA-dependent RNA polymerase and the major capsid protein VP60) and ORF2 encoding the VP10 protein a minor structural protein^9,10^.

RHD was first reported in China in 1984^11^: where over 140 million rabbits were killed in the course of an outbreak^12^. Subsequent spread of the virus was reported in Europe and other continents^13–19^. In Africa, RHDV has been reported since the late 1980s^20^ but molecular characterization and diversity of the circulating variants and their impact are yet to be understood.

There are six serogroups (G1-G6) RHDV^21^ and one serotype^22^ before 2010. These groups are known as “classical RHDV”. A new strain RHDV2 (GI.2 or RHDVb) was discovered in France in 2010^23^ and is now reported in many parts of Europe^22, 24–26^ and other parts of the world^27, 28^. RHDV2 was responsible for major outbreaks causing deaths in previously vaccinated adult rabbits as well as younger rabbits known to be immune to classical RHD^22, 23, 29^. This strain is now reported in Europe, Australia, Canada and more recently the USA^27, 30, 31^. RHD has been reported in some African countries like Tunisia, Morocco, Cape Verde and Egypt^7, 32–34^. In sub-Saharan Africa, outbreaks of GI.2 were reported and confirmed by enzyme immunoassays (EIA) in domestic rabbits from Benin Republic^34^, and PCR from Cote d’Ivoire^30^. However, to date there are no available virus genome data from any Sub-Saharan African country, rendering it difficult to trace their origin, evolution and genetic diversity.

Following devastating outbreaks of suspected cases of RHD affecting several farmers from the southwestern region of Nigeria, there was considerable loss of investment with no surviving rabbits in some of the establishments. During the decline in the number of cases in August 2020, two farms (Farm A and Farm B) reported high mortalities and were visited to carry out post-mortem examinations and sampling for molecular diagnosis. During the outbreak, farmers had observed symptoms similar to those caused by RHDV. The fatalities affected exotic breeds of rabbits of all ages and both sexes. The symptoms reported by farmers were anorexia (a day prior to death), lacrimal discharge, lethargy, bleeding from the oral and nasal orifices and sudden death.

Here we report the use of microbial metagenomics sequencing to uncover the first genomic characterization and whole genome sequencing of the RHDV2 responsible for an outbreak of RHD in Sub-Saharan Africa.

## Results

### Gross Post mortem findings

The carcasses were found to be in good body condition and slightly autolyzed (5/5). The oral and ocular mucous membranes were mildly pale (5/5). There were a few multifocal widespread petechial haemorrhages on the ventral and dorsal abdominal subcutaneous muscle (2/5), as well a few multifocal petechial haemorrhages on the pleura surface of the lungs (2/5).

### RT-PCR

Two samples (RT2 and RT4) out of five were positive by RT-PCR for RHDV targeting the VP60 gene (Table 1). Bands were confirmed on electrophoresis at regions between 1500 and 2000bp (Results not shown).

**Table 1:** Sample Demographics

| Sample ID | Sex | Age | RT-PCR | Genome Sequence |
| --- | --- | --- | --- | --- |
| RT1 | Female | 8 weeks | Negative | No |
| RT2 | Male | 8 weeks | Positive | Yes (partial) |
| RT3 | Male | 5 weeks | Negative | No |
| RT4 | Male | 3 weeks | Positive | Yes (partial) |
| RT5 | Male | Adult | Negative | Yes (full) |

### Sequencing

We obtained one full genome (sample RT5) and two partial sequences (samples RT4 and RT2) (Table 1). Metagenomics analysis identified the presence of Rabbit haemorrhagic disease virus (RHDV) reads in three samples (RT2, RT4 and RT5) out of five samples sequenced. Analysis of the sequence data revealed that all three of our sequences belong to genotype RHDV2 also known as GI.2 or RHDVb. BLAST analysis revealed a sample RT5 shared a 98.58% identity with a RHDV2 strain from the Netherlands, RT4 shared a 99.05% identity with a RHDV2 strain from while RT2 shared a 100% with the same RHDV2 strain from Germany as RT4 (Figure 1). Phylogenetic analysis further confirmed the results of our BLAST analysis, demonstrating that the sequences from these animals belong to the RHDV2 genotype as they clustered together in the same clade with previous RHDV2 sequences from Europe and other parts of Africa (Figure 1). Analysis of Single nucleotide polymorphisms (SNPs) in the RHDV2 genome of sample RT5 (which was negative by RT-PCR) obtained from our study, revealed 11 unique mutations resulting in amino acid changes. These are: Thr179Ile, Leu216Ser, Leu342Ser, Arg827Lys, Gln928Arg, Thr1337Ile, Thr1742Ile, Leu1929Pro, Ser1978Phe, Val2104Ala, Val2127Ala. Four of these mutations (Leu1929Pro, Ser1978Phe, Val2104Ala, Val2127Ala) occur in the region of target for the RT-PCR primers. Sample RT4 had a unique mutation Pro440Leu compared to other RHDV2 sequences they clustered with on the tree.

**Figure 1:**
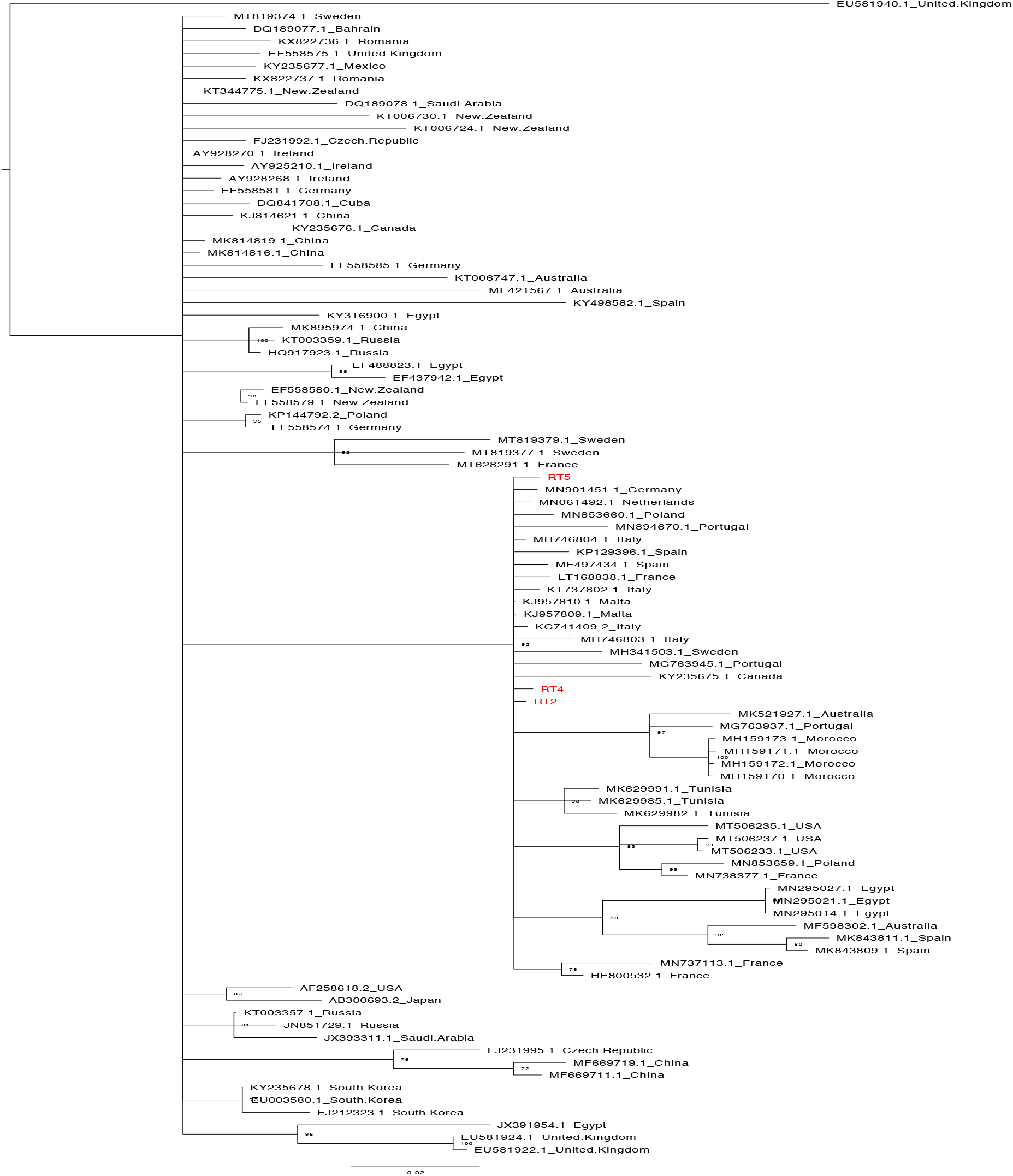
Phylogenetic relationship between the sequences from this study (coloured red) and representative RHDV sequences obtained from the NCBI database

## Discussion

Our initial suspicion of RHD based on case history, the extent of morbidity and mortality and post mortem findings was confirmed by RT-PCR in two out of five tissue samples. Further metagenomic sequence analysis resulted in three genome (one full and two partial) assemblies.

To the best of our knowledge, this study represents the first full genome sequence of Rabbit Hemorrhagic Disease Virus (RHDV) in sub-Saharan Africa. Bioinformatics analysis of the virus genomes revealed that the three sequences from this study belong to genotype *Lagovirus* europaeus/GI.2 also known as RDHV2 or RHDVb. This genotype was first identified in 2010 as a novel pathogenic form of *Lagovirus* in France^8^ after which it spread rapidly through Europe and other parts of the world (Oceania, North America and Africa)^27, 28, 30–33^. It has also been reported to replace the former circulating GI.1 strain in Australia and Portugal^35, 36^. Phylogenetic analysis showed that our sequences clustered closely with previous sequences from Europe. BLAST analysis revealed that sample RT5 shared a 98.58% identity with a RHDV2 strain from the Netherlands, RT4 shared a 99.05% identity with a RHDV2 strain from Germany while RT2 shared a 100% with the same RHDV2 strain from Germany as RT4 suggesting that the virus was most likely imported into the country from Europe.

The *Lagovirus* europaeus/GI.2 strain is known to dominate in most regions it has been found and most recently in Morocco^37^. RHDV has been circulating in some African countries and more recently, there have been reports of GI.2 outbreaks in a few countries from North Africa (Tunisia, Egypt and Morocco)^37^.

RHDV strains have yet to be characterized in Sub-Saharan African despite its devastating effect on rabbit farms and the industry. Given the growing distribution and the devastating effect, there is a need for heightened surveillance. In Australia, it spread to all states and territories and rapidly became the dominant circulating strain within 18 months of initial detection^36^. Ongoing surveillance and genomic sequencing are therefore needed to understand the interaction of the various *Lagoviruses* in Nigeria and their impact on host populations. Next generation sequencing of more samples is also needed in order to determine the time of introduction of the virus into the country and how it is spreading through the country.

Our phylogenetic analysis also revealed high genetic diversity of the GI.2 strain as shown by the number of branching events within the clade. In addition, the high number of unique mutations observed in the sequences from this study further emphasizes this diversity, which is likely due in part to the rapid spread and evolution of this virus. Also, it may explain why it affects rabbits of all ages more or less equally in our communities. The fact that some of the mutations occurred in the region of target of the RT-PCR primers may be the reason why some of the RT-PCR were negative (RT5), and yet yielded a full genome after NGS and metagenomic analysis. This finding emphasizes the need to develop primers for diagnosis based on RHDV sequences from the country of outbreak in order to avoid false negative RT-PCR results. Compiled genomic data should be carefully considered when developing diagnostics and vaccines.

The findings from this study is a significant landmark in the field, as it has revealed the circulation of the RHDV2 in Nigeria and reports the first genomics characterization of RHDV in sub-Saharan Africa. The close sequence homology suggests that the virus was most likely imported from Europe. In addition, the high genetic diversity of the GI.2 strains found highlights the need for characterization of many more samples across sub-Saharan Africa and in high incidence zones to guide the development of improved diagnostics and vaccines.

The detection of RHDV2 on unbiased metagenomic sequencing, as shown in this study illustrates the power of genomics in explaining a suspected outbreak. This ability to rapidly identify and characterize an emerging virus (RHDV 2) – in an unusual cluster– highlights the value of in-country genomics capacity. Serology using ELISA and RT-PCR are the current methods of choice for RHDV diagnosis in Sub-Saharan Africa. These diagnostic methods despite their limitations are done in very few selected laboratories. The integration of genomics capacity into the established, but siloed, pathogen-specific diagnostic platforms provides exciting opportunities for Veterinary public health surveillance.

## METHODS

### Post Mortem and Sample collection

In August 2020, following reports of devastating outbreaks of suspected Rabbit Haemorrhagic Disease (RHD) in Rabbit farms around, Ibadan, South-western region of Nigeria, post mortem was carried out on four carcasses from farm A and one carcass from farm B. Signalment from farm A consists of 1 female (8 weeks old) and 3 males (8 weeks, 5 weeks and 3weeks old). The breeds on the farm were Hyla. Farm B consists of one adult male Hyla. Tissue sections were collected into RNAlater from the five (5) rabbit carcasses suspected to have died from RHD. The samples were tagged RT1, RT2, RT3, RT4 (farm A) and RT5 (farm B). Tissues (liver, spleen, lungs) of each animal were pooled for RT1 - RT4 while only liver was collected for RT5. The samples were then maintained in a cold chain and RNALater during transportation to the African Centre of Excellence for the Genomics of Infectious Disease (ACEGID), Redeemer’s University, Ede, Nigeria for PCR confirmation and metagenomics sequencing analysis.

### RNA Extraction and RT-PCR

Samples were first washed in PBS and thereafter homogenized and macerated. Total RNA was extracted from tissues macerated in TRIzol using QIAamp Viral RNA extraction kit (Qiagen, Hilden, Germany) according to manufacturer’s instructions. Extracted RNA was stored in −20°C until RT-PCR and sequencing.

On the 16^th^ of September, 2020, RT-PCR was performed on extracted RNA based on modified established protocol^14^ with primers targeting the VP60 gene. One-step SuperScript™ III One-Step RT-PCR System with Platinum™ Taq DNA Polymerase (Invitrogen, USA) was used for PCR amplification targeting the VP60 capsid protein giving a 1740 base pair product^11^. The RT-PCR final reaction volume of 25ul was made up of 12.5 μl of 2X reaction mix, 1.25μl of 20μM each of forward and reverse primers, 1μl SuperScript™ III RT/Platinum™ Taq Mix, 1μl RNA template and MgSo4 optimization to a final concentration of 2.5μM and nuclease free water to make up the reaction volume. Our thermocycling condition includes; 55 °C for 30 min, 95 °C for 15 min, then 40 cycles at 95 °C for 1 min, 58 °C for 30 s and 72 °C for 1 min, and a final extension step at 72 °C for 10 min. PCR products were viewed in 1% gel electrophoresis (Results not shown).

### Next Generation Sequencing and Bioinformatics Analysis

Upon RT-PCR confirmation on September 25, 2020, Nextera XT sequencing Libraries were made based on established unbiased protocol^38^. Briefly, RNA was turbo DNase treated to remove contaminating DNA, and converted to cDNA. Nextera XT sequencing libraries were made from cDNA. Libraries were sequenced using the Miseq Illumina platform in our sequencing platform at ACEGID, Redeemer’s University, Ede, Osun State, Nigeria.

Following sequencing, raw reads from the next-generation sequencing machine were uploaded to our cloud-based platform (DNAnexus, www.dnanexus.com). Quality control was carried out on the raw reads using fastqc (https://www.bioinformatics.babraham.ac.uk/projects/fastqc). Metagenomics analysis was carried out using Kraken2^39^.

RHDV genomes were assembled using our publicly-available software viral-ngs v2.1.8 (https://github.com/broadinstitute/viral-ngs) implemented on DNAnexus. Following BLASTn analysis, ninety representative RHDV sequences were obtained from the NCBI database and aligned with our three sequences using MAFFT v7.388^40^ with further adjustment made manually as necessary in Geneious^41^. A neighbour-joining tree was generated also in Geneious from a distance matrix corrected for nucleotide substitutions by the Tamura-Nei model. The tree was viewed and manually edited using FigTree (http://tree.bio.ed.ac.uk/software/figtree/). We aligned the three sequences from this study with all sequences in the same RHDV2 clade on the tree to check for amino acid mutations specific to our new sequences from Nigeria. We also mapped the RT-PCR primers to our full genome obtained from this study to check if any mutation observed occurred in the target regions of the primers.

## Data Availability

All sequences from this study were submitted to the National Center for Biotechnology Information (NCBI) database and the accession numbers (MW123059 - MW123061) received on the 16 October, 2020.

## Acknowledgements

This work is supported by the BBSRC-OVEL project BB/R020116/1 and WELLCOME TRUST project 216619/Z/19/Z to CTH and JLH. We are grateful to Mrs. Philomena Eromon (African Centre of Excellence for Genomics of Infectious Diseases (ACEGID), Redeemer’s University, Ede, Osun State, Nigeria) for advice during the RT-PCR trouble-shooting.

## Authors Contribution

A.N.H.-Initiated the work, performed the necropsy examination, advised on NGS analysis and co wrote the first draft. O.A.O.-performed the necropsy examination and sample collection, performed RT-PCR and analysis, and co-wrote the first draft. J.U.O. - performed the RT-PCR and NGS experiments and co-wrote the first draft. P.E.O. - performed Bioinformatics analysis of NGS data and discussed them. A.S.O.-participated in the necropsy examination and sample collection, revised the first draft. J. L. H.-Reviewed the manuscript. C.T.H.-Guided the NGS experiments, genomic data analysis and discussions, review the manuscript. All authors contributed to the final version of the manuscript.

## Additional Information

No additional information available

## References

1. Meyers, G., Wirblich, C., & Thiel, H. J. Rabbit hemorrhagic disease virus-molecular cloning and nucleotide sequencing of a calicivirus genome. Virology 184, 664–676. https://doi:10.1016/0042-6822(91)90436-F (1991).

2. Guittre, C. I., Baginski, G. & Gall, L. E. Detection of rabbit hemorrhagic disease virus isolates and sequence comparison of the N-terminus of the capsid protein gene by the polymerase chain reaction. Res. Vet. Sci. 58, 128–132 (1995).

3. Abrantes, J., van der Loo, W., Le Pendu, J. & Esteves, P. J. Rabbit haemorrhagic disease (RHD) and rabbit haemorrhagic disease virus (RHDV): a review. Vet. Res. 43, 12. https://doi.org/10.1186/1297-9716-43-12 (2012).

4. Mitro, S. & Krauss, H. Rabbit hemorrhagic disease: A review with special reference to its epizootiology. Eur. J. Epidemiol. 9, 70–78 (1993).

5. Asgari, S., Hardy, J. R., Sinclair, R. G. & Cooke, B. D. Field evidence for mechanical transmission of rabbit haemorrhagic disease virus (RHDV) by flies (*Diptera Calliphoridae*) among wild rabbits in Australia. Virus Res. 54(2) 123–132 (1998).

6. Ohlinger, V. F., Haas, B. & Thiel, H. J. Rabbit haemorrhagic disease (RHD): Characterization of the causative calicivirus. Vet. Res. 24(2):103–116 (1993).

7. World Organization for Animal Health (OIE) 2010. Terrestrial manual. In:Rabbit Haemorrhagic Disease. 2010 Ch. 2.6.2. OIE Paris. France. https://www.oie.int/fileadmin/Home/eng/Animal_Health_in_the_World/docs/pdf/Disease_cards/RHD.pdf (2010).

8. Le Gall-Reculé, G. et al. Detection of a new variant of rabbit haemorrhagic disease virus in France. Vet. Rec. 168, 137–138 (2011)

9. Dalton, K. P. et al. Vaccine breaks: Outbreaks of myxomatosis on Spanish commercial rabbit farms. Vet. Microbiol. 178, 208–216 (2015).

10. Meyers, G., Wirblich, C., Thiel, H. J. & Thumfart, J. O. Rabbit hemorrhagic disease virus: genome organization and polyprotein processing of a calicivirus studied after transient expression of cDNA constructs. Virology 276, 349–363 (2000).

11. Liu, S. J., Xue, H. P., Pu, B. Q., Qian, N. H. & Others. A new viral disease in rabbits. Zhongguo Xu Mu Shou Yi 16, 253–255 (1984).

12. Xu, W. Y. Viral haemorrhagic disease of rabbits in the People’s Republic of China: epidemiology and virus characterisation. Rev. Sci. Tech. 10, 393–408 (1991).

13. Rahali, N., Sghaier, S., Kbaier, H., Zanati, A. & Bahloul, C. Genetic characterization and phylogenetic analysis of rabbit hemorrhagic disease virus isolated in Tunisia from 2015 to 2018. Arch. Virol. 164, 2327–2332. https://doi:10.1007/s00705-019-04311-z (2019).

14. Ismail, M. M., Mohamed, M. H. A., El-Sabagh, I. M. & Al-Hammadi, M. A. Emergence of new virulent rabbit hemorrhagic disease virus strains in Saudi Arabia. Trop. Anim. Health Prod. 49, 295–301. https://doi:10.1007/s11250-016-1192-5 (2017).

15. Gould, E. A. First case of rabbit haemorrhagic disease in Canada: contaminated flying insect, vs. long-term infection hypothesis: NEWS AND VIEWS: REPLY. Mol. Ecol. 21, 1042–1047. https://doi:10.1111/j.1365-294X.2012.05462.x (2012).

16. Fitzner, A. & Niedbalski, W. Phylogenetic analysis of rabbit haemorrhagic disease virus (RHDV) strains isolated in Poland. Arch. Virol. 162, 3197–3203. https://doi:10.1007/s00705-017-3476-0 (2017).

17. Elliott, S. & Saunders, R. Rabbit haemorrhagic disease in the UK. Vet. Rec. 181, 516–516. https://doi:10.1136/vr.j5149 (2017).

18. Bergin, I. L. et al. Novel calicivirus identified in rabbits, Michigan, USA. Emerg. Infect. Dis. 15, 1955–1962. https://doi:10.3201/eid1512.090839 (2009).

19. Abrantes, J. et al. New variant of rabbit hemorrhagic disease virus, Portugal, 2012-2013. Emerg. Infect. Dis. 19, 1900–1902. https://doi:10.3201/eid1911.130908 (2013).

20. Morisse, J. P., Le Gall, G. & Boilletot, E. Hepatitis of viral origin in Leporidae: introduction and aetiological hypotheses. Rev. Sci. Tech. 10, 269–310 (1991).

21. Le Gall-Reculé, G. et al. Phylogenetic analysis of rabbit haemorrhagic disease virus in France between 1993 and 2000, and the characterisation of RHDV antigenic variants. Arch. Virol. 148, 65–81 (2003).

22. Dalton, K. P. et al. Variant rabbit hemorrhagic disease virus in young rabbits, Spain. Emerg. Infect. Dis. 18, 2009–2012 (2012).

23. Carvalho, C. L. et al. Emergence of rabbit haemorrhagic disease virus 2 in the archipelago of Madeira, Portugal (2016-2017). Virus Genes 53, 922–926. https://doi:10.1007/s11262-017-1483-6 (2017).

24. Baily, J. L., Dagleish, M. P., Graham, M., Maley, M. & Rocchi, M. S. RHDV variant 2 presence detected in Scotland. Vet. Rec. 174, 411. doi: 10.1136/vr.g2781 (2014).

25. Westcott, D. G. & Choudhury, B. Rabbit haemorrhagic disease virus 2-like variant in Great Britain. Vet. Rec. 176, 74. https://doi:10.1136/vr.102830 (2015).

26. Hall, R. N. et al. Emerging Rabbit Hemorrhagic Disease Virus 2 (RHDVb), Australia. Emerg. Infect. Dis. 21, 2276–2278 (2015).

27. Martin-Alonso, A., Martin-Carrillo, N., Garcia-Livia, K., Valladares, B. & Foronda, P. Emerging rabbit haemorrhagic disease virus 2 (RHDV2) at the gates of the African continent. Infect. Genet. Evol. 44, 46–50. https://doi:10.1016/j.meegid.2016.06.034 (2016).

28. Dalton, K. P., Nicieza, I., Abrantes, J., Esteves, P. J. & Parra, F. Spread of new variant RHDV in domestic rabbits on the Iberian Peninsula. Vet. Microbiol. 169, 67–73 (2014).

29. World Organization for Animal Health (OIE) 2016 Rabbit haemorrhagic disease CoteD’Ivoire https://www.oie.int/wahis_2/public/wahid.php/Reviewreport/Review?page_refer=MapFullEventReport&reportid=20852 (2016).

30. World Organization for Animal Health (OIE) 2020. Rabbit Haemorrhagic Disease. https://www.aphis.usda.gov/publications/animal_health/fs-rhdv2.pdf (2020).

31. World Organization for Animal Health (OIE) 2005. World Animal Health Publication and Handistatus II (data set for 2004). Paris, France:Office International des Epizooties. https://www.oie.int/animal-health-in-the-world/the-world-animal-health-information-system/data-before-2005-handistatus/ (2005).

32. World Organization for Animal Health (OIE) 2009. World Animal Health Information Database - Version: 1.4. World Animal Health Information Database. Paris, France: World Organisation for Animal Health. http://www.oie.int (2009).

33. World Organization for Animal Health (OIE) 2015. Rabbit haemorrhagic disease, Benin. https://www.oie.int/wahis_2/public/wahid.php/Reviewreport/Review?page_refer=MapEventSummary&reportid=19903 (2015).

34. Erfan, A. M. & Shalaby, A. G. Genotyping of rabbit hemorrhagic disease virus detected in diseased rabbits in Egyptian Provinces by VP60 sequencing. Vet World 13, 1098–1107 (2020).

35. Lopes, A. M., Correia, J., Abrantes, J., Melo, P. & Ramada, M. Is the new variant RHDV replacing genogroup 1 in Portuguese wild rabbit populations? Viruses 7, 27–36 (2015).

36. Mahar, J. E. et al. Rabbit Hemorrhagic Disease Virus 2 (RHDV2; GI.2) Is Replacing Endemic Strains of RHDV in the Australian Landscape within 18 Months of Its Arrival. J. Virol. 92, 01374–17. https://doi.org/10.1128/JVI.01374-17 (2018).

37. Lopes, A. M., Rouco, C., Esteves, P. J. & Abrantes, J. GI.1b/GI.1b/GI.2 recombinant rabbit hemorrhagic disease virus 2 (Lagovirus europaeus/GI.2) in Morocco, Africa. Arch. Virol. 164, 279–283. https://doi.org/10.1007/s00705-018-4052-y (2019).

38. Matranga, C. B. et al. Unbiased Deep Sequencing of RNA Viruses from Clinical Samples. J. Vis. Exp. 113, 54117. https://doi.org/10.3791/54117 (2016).

39. Wood, D. E., Lu, J. & Langmead, B. Improved metagenomic analysis with Kraken 2. Genome Biol. 20, 257. https://doi.org/10.1186/s13059-019-1891-0 (2019).

40. Katoh, K., Misawa, K., Kuma, K.-I. & Miyata, T. MAFFT: a novel method for rapid multiple sequence alignment based on fast Fourier transform. Nucleic Acids Res. 30, 3059–3066 (2002).

41. Kearse, M. et al. Geneious Basic: an integrated and extendable desktop software platform for the organization and analysis of sequence data. Bioinformatics 28, 1647–1649 (2012).

